# Uninterpretable interactions: epistasis as uncertainty

**DOI:** 10.1101/378489

**Authors:** Zachary R. Sailer, Michael J. Harms

## Abstract

Epistasis is a common feature of genotype-phenotype maps. Understanding the patterns of epistasis is critical for predicting unmeasured phenotypes, explaining evolutionary trajectories, and for inferring the biological mechanisms that determine a map. One common approach is to use a linear model to decompose epistasis into specific pairwise and high-order interactions between mutations. Such interactions are then used to identify important biology or to explain how the genotype-phenotype map shapes evolution. Here we show that the coefficients extracted from such analyses are likely uninterpretable. They cannot be extracted reliably from experimental genotype-phenotype maps due to regression bias. Further, we can generate epistatic “interactions” indistinguishable from those in experimental maps using a completely random process. From this, we conclude that epistasis should be treated as a random, but quantifiable, variation in these maps. This perspective allows us to build predictive models with known error from a small number of measured phenotypes. It also suggests that we need mechanistic, nonlinear models to account for epistasis and decompose genotype-phenotype maps.

## Introduction

Epistasis—that is, non-additivity between mutations—is a ubiquitous feature of genotype-phenotype maps (Fowler et al., 2010; Weinreich, 2011; Weinreich et al., 2013; Yokoyama et al., 2014; Anderson et al., 2015; Palmer et al., 2015; Podgornaia and Laub, 2015; Doud and Bloom, 2016; Boyle et al., 2017; Hopf et al., 2017; Chan et al., 2017; Sailer and Harms, 2017a; Starr et al., 2017; Domingo et al., 2018; Weinreich et al., 2018). Epistasis can provide mechanistic insight into the determinants of phenotypes (Schreiber and Fersht, 1995; Horovitz, 1996; Ritchie et al., 2001; Segrè et al., 2005); however, it also complicates predicting unmeasured phenotypes (de Visser and Krug, 2014; Miton and Tokuriki, 2016; Sailer and Harms, 2017b; Nyerges et al., 2018), as the effect of a mutation changes depending on the presence or absence of other mutations. Despite a century of work (Fisher, 1918), epistasis remains challenging to analyze and interpret (Cordell, 2002; Phillips, 2008; Crow, 2010; Weinreich et al., 2013, 2018).

One approach is to decompose epistasis into specific pairwise and high-order interactions between mutations (Heckendorn and Whitley, 1999; Weinreich et al., 2013; Poelwijk et al., 2016; Sailer and Harms, 2017a; Poelwijk et al., 2017; Weinreich et al., 2018). This is often done by treating each coefficient as a linear and independent perturbation to the additive phenotype (Heckendorn and Whitley, 1999; Poelwijk et al., 2016). Such an approach is a direct extension of classic approaches in quantitative genetics and biochemistry. In a genetics context, one might measure the effect of a mutation in two genetic backgrounds to dissect metabolic and regulatory pathways (Ritchie et al., 2001; Segrè et al., 2005). Likewise, mutant cycles are a mainstay of biochemistry. Introducing mutations individually and together allows one to infer the nature of physical interactions between residues in macromolecules (Schreiber and Fersht, 1995; Horovitz, 1996).

Although linear epistasis models are very commonly used (Weinreich et al., 2013; Yokoyama et al., 2014; Anderson et al., 2015; Palmer et al., 2015; Starr et al., 2017; Domingo et al., 2018), two recent observations raise questions about their utility. The first is that regression can lead to biased estimates of linear epistatic coefficients, and thus poor predictive power of epistatic models (Otwinowski and Plotkin, 2014). The second is that one can generate maps with extensive pairwise and high-order epistasis using a toy model of proteins that do not explicitly include such interactions (Sailer and Harms, 2017b). This indicates that there may be no simple way to relate linear epistatic coefficients back to underlying biology, thus undermining their utility as indicators of biological mechanism.

Motivated by these concerns, we set out to systematically investigate linear epistatic models constructed from twelve published genotype-phenotype maps. We focused on two criteria for utility: the ability of such models to predict unmeasured phenotypes and the ability of such coefficients to provide mechanistic insight into the map. We studied maps for which all 2^*L*^ combinations of *L* mutations were measured. Because these maps have the same number of observations as coefficients in a high-order epistatic model, they can be readily decomposed into epistatic coefficients from second to *L^th^*-order. Further, the selected maps cover many different classes of genotypes, phenotypes, and total magnitudes of epistasis.

We find that the epistatic coefficients we extract by regression from such maps are quite poor at predicting unmeasured phenotypes. This arises from bias in the regressed coefficients—exactly as predicted by Otwinowski and Plotkin (Otwinowski and Plotkin, 2014). Further, we find we can generate epistatic coefficients similar to experimental coefficients by simply using randomly assigned phenotypes. This suggests that the pairwise and high-order interactions we extract are likely decompositions of random noise. We therefore propose that we should not decompose genotype-phenotype maps into specific interactions between mutations using linear models. Rather, in the context of a whole genotype-phenotype map, epistasis is best interpreted as a global metric capturing roughness (Szendro et al., 2013; Ferretti et al., 2018). This translates directly to a measure of uncertainty on predicted phenotypes, as well as an indication that an improved mechanistic model is required.

## Materials and Methods

### Linear epistasis models

We used a linear epistasis model to decompose genotype-phenotype maps into up to *L^th^*-order epistatic coefficients. The model is linear in that it consists of a collection of independent epistatic coefficients that are summed to describe each phenotype (Fisher, 1918; Poelwijk et al., 2016). (The assumption of linearity contrasts with other models, such as a Potts model, in which mutations sum in a nonlinear fashion (Hopf et al., 2017)). There are two common formulations a linear epistasis model, the Hadamard model (sometimes called a Walsh or Fourier model) and the biochemical model (Poelwijk et al., 2016). The approaches differ in their choice of coordinate origin. Each model has been described in detail elsewhere (Heckendorn and Whitley, 1999; Weinreich et al., 2013; Poelwijk et al., 2016). The two models are related by a simple set of linear transformations (Poelwijk et al., 2016). Throughout the text, we describe our results using the Hadamard model, but our conclusions are robust to the choice of model (see supplemental figures referenced throughout the text).

The Hadamard model uses the geometric center of the map as the coordinate origin (Heckendorn and Whitley, 1999; Weinreich et al., 2013; Poelwijk et al., 2016; Sailer and Harms, 2017a). Each genotype is made up of *L* sites. In a binary genotype-phenotype map, the sites have two possible states: “wildtype” or “derived”. Both states have equal effects but opposite signs. Each mutation is treated as a linear perturbation away from the origin of the map,

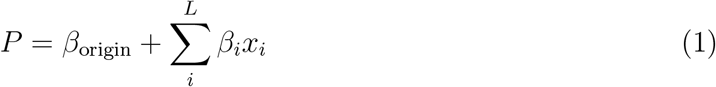

where *β*_origin_ is the origin of the genotype-phenotype map, *β_i_* is the effect of site *i*, and *x_i_* is 1 if site *i* is “wildtype” and – 1 if “derived”. We can then add linear coefficients to describe interactions between mutations to Eq. 1. For pairwise interactions, this has the form:

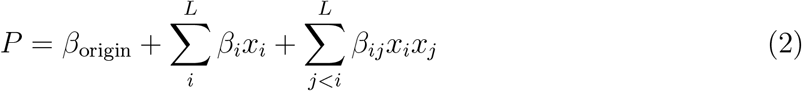

where *β_ij_* is a pairwise epistatic coefficient. For the high-order model, the expansion continues:

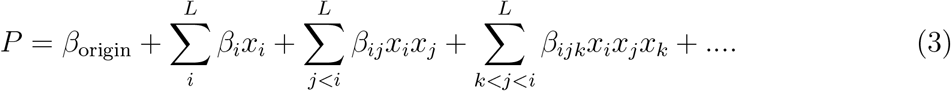

The model can be expanded all the way to *L^th^*-order interactions.

### Linearizing experimental genotype-phenotype maps

Prior to extracting epistatic coefficients from experimental genotype-phenotype maps, we corrected each map for *global* epistasis, which arises when mutations combine on some scale other than an additive scale (Chou et al., 2011; Tokuriki et al., 2012; Schenk et al., 2013; Sailer and Harms, 2017a; Otwinowski et al., 2018). This violates the assumption of linearity inherent in the epistasis models (Fisher, 1918; Cordell, 2002; Sailer and Harms, 2017a). Global epistasis manifests as a non-normal distribution of the residuals between the 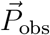 (the vector of observed phenotypes) and 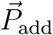 (the vector phenotypes calculated using an additive model) (Sailer and Harms, 2017a; Otwinowski et al., 2018). Such epistasis can be minimized by identifying a nonlinear function *T* that captures global curvature in the relationship between 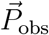 and 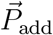, yielding normally distributed fit residuals (Box and Cox, 1964; Szendro et al., 2013; Sailer and Harms, 2017a; Otwinowski et al., 2018):

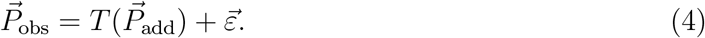

where 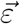 are the fit residuals. We linearized all experimental maps by fitting a second-order spline to the 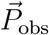 vs. 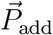 curve for each map prior to extracting linear epistatic coefficients (Otwinowski et al., 2018).

### Epistasis and linear regression

We used linear regression to regress epistasis models against experimental and simulated genotype-phenotype maps. We formulated the problem as follows:

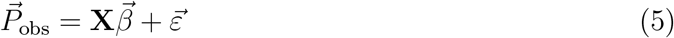

where 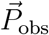 is a vector of observed phenotypes (corrected for global epistasis), 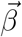 is a vector of epistatic coefficients, **X** is a matrix that encodes the sign of each coefficient according to Eq. 3, and 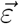 is a vector of residuals. The goal was to estimate coefficients in 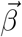 that minimized the magnitudes of the values in 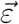.

We used three different regression approaches: ordinary least-squares, lasso, and ridge. The number of coefficients in these maps grows rapidly with the number of sites. For a binary map with *L* sites, there are 2^*L*^ possible fit coefficients. Lasso and ridge regression are strategies to identify only those coefficients that contribute significantly to the variation in the data. These strategies have been used previously to dissect linear epistatic models (Otwinowski and Plotkin, 2014; Poelwijk et al., 2017). Throughout the text, we describe results using lasso regression, but our conclusions are robust to the choice of regression strategy (see supplemental figures referenced throughout the text).

### Simulating epistatic genotype-phenotype maps

We constructed genotype-phenotype maps using Equations 1 and 3. First, we set the additive coefficients to random values drawn from a normal distribution. We then added all 2^*nd*^-through *L^th^*-order epistatic coefficients. We set the values of the coefficients to random values drawn from a different normal distribution. The widths of the additive and epistatic distributions were tuned to match the relative magnitudes of epistatic coefficients extracted from experimental maps. Further, we could tune the fraction of epistasis in a simulated genotype-phenotype map by changing the relative widths of the additive and epistatic distributions with respect to one another.

### Software

We implemented the epistasis models using Python 3 extended with *numpy, scipy*, and *pandas* (van der Walt et al., 2011; McKinney, 2010). We used the Python package *scikit-learn* to perform ordinary-, lasso-, and ridge-regression (Pedregosa et al., 2011). We used the Python package *Imfit* to perform nonlinear-least squares regression (Newville et al., 2018). Plots were generated using *matplotlib* and *Jupyter* notebooks (Hunter, 2007; Perez and Granger, 2007). Our full software packages are available in the *gpmap* (https://harmslab.github.com/gpmap) and *epistasis* (https://harmslab.github.com/epistasis) packages on Github.

### Data availability statement

All software is available for download from:

- https://github.com/harmslab/gpmap
- https://github.com/harmslab/epistasis.

Data sets are available from:

- https://github.com/harmslab/genotype-phenotype-maps.

Supplemental figures S1-S4 are available via GSA Figshare.

## Results

### Regression yields biased estimates of epistatic coefficients

We started with a straightforward question: What fraction of a genotype-phenotype map must we observe to resolve a linear epistatic model that predicts unmeasured phenotypes? We simulated a genotype-phenotype map consisting of all 2^8^ binary combinations of 8 mutations. We then assigned random epistatic coefficients using an 8th-order Hadamard matrix, such that epistasis accounted for 20% of the variation in phenotype (see methods). The epistatic coefficients were similar in magnitude and sign to those extracted from experimental genotype-phenotype maps (Fig S1).

To test our ability to predict phenotypes, we masked a fraction of the genotypes, fit linear epistatic models to the unmasked genotypes, and attempted to predict the masked genotypes.

We then calculated the correlation between the model and unmasked observations 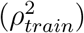 and the model and masked observations 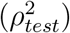. We repeated this for 1,000 pseudo-replicate training and test sets.

As a starting point, we fit the additive model (Eq. 1). We found that the additive model converged on 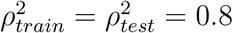 when 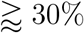 of the map was used for the fit (red lines, Fig 1A). The model converges once each mutation has been observed across a sufficient number of genetic backgrounds to average out the epistatic perturbations to the phenotype. Because, by construction, 20% of the variation in the map is due to epistasis, the best the additive model can do is explain 80% of the variation in phenotype.

**Figure 1:**
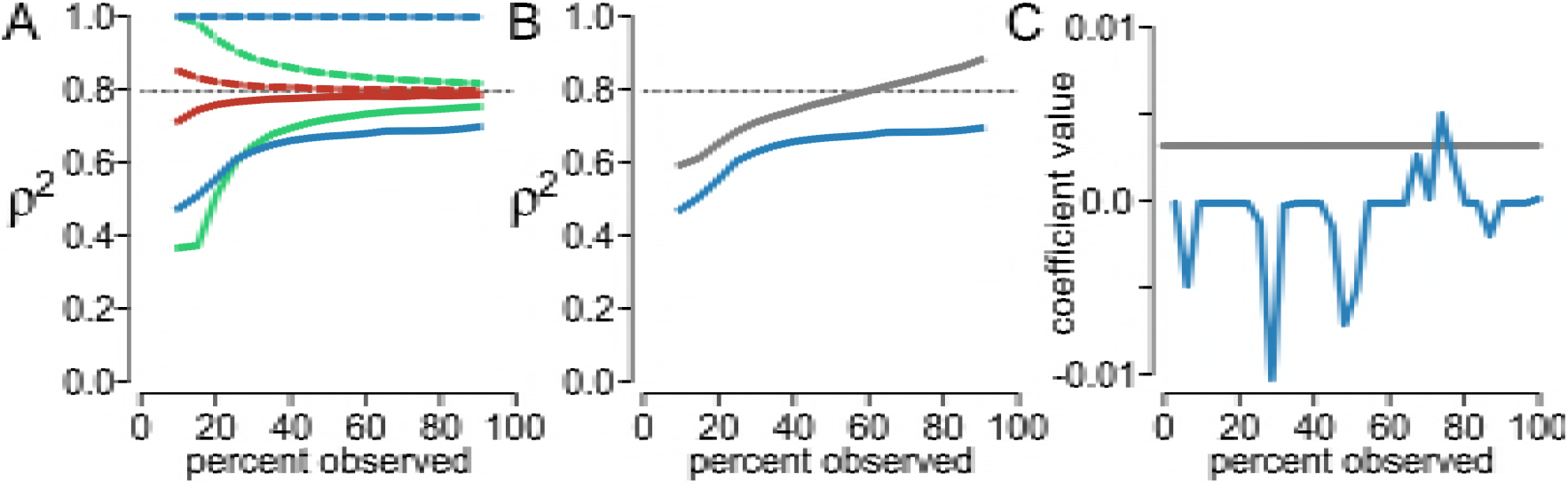
Linear epistatic coefficients cannot be estimated from an incomplete, simulated genotype-phenotype map. A) Fit scores versus the percent of the genotypes in the map used to train the model, from 10% to 90%. The dashed gray line indicates the amount of additive variation in the map (80%). Colors indicate model order: additive (red), pairwise epistasis (green), and high-order epistasis (blue). Dashed lines indicate 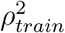 and solid lines indicate 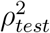. B) Fit scores versus the fraction of map used to train the model. Blue curve uses regressed coefficients (reproduced from panel A). Gray curve shows 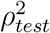 if we use the coefficients used to generate the map. C) Value of a pairwise epistatic coefficient as we add data to the fit. Gray line indicates the value of the coefficient used to generate the map.

We next tried to improve our predictive power by adding coefficients describing either pairwise interactions between mutations (Eq. 2) or all interactions (up to eighth-order) (Eq. 3). Because Eq. 3 is the model we used to generate the map, this model should, in principle, be able to explain all variation in the map.

We found that neither the pairwise nor high-order models performed as well as the additive model (green and blue lines, Fig 1A). Even when 90% of the genotypes were included in the training set, the pairwise and high-order models had 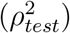 of 0.73 and 0.62—much less than the value of 0.80 achieved by the additive model. Worse, this failure to predict the test set was accompanied by much higher 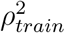 values. The high-order model, in particular, had a correlation of 1.0 with the training set (Fig 1A, dashed blue line), even while test set correlation languished around 0.6 (Fig 1A, solid blue line).

This result arises because regression yields biased estimates of the epistatic coefficients (Otwinowski and Plotkin, 2014). We know that high-order epistasis is present, because we used the same high-order model we are now fitting to generate the underlying map. The fit coefficients, however, do not accurately capture this variation. This can be seen in Fig 1B. The blue line reproduces 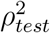 for the high-order model from Fig 1A. The gray curve shows values of 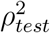 calculated using the epistatic coefficients used to generate the map. The divergence between these curves indicates that the regression fails to extract the correct values for the epistatic coefficients.

This can also be seen by examining the values of the extracted epistatic coefficients. The blue curve in Fig 1C shows the estimated value of a single, pairwise epistatic coefficient within the high-order model as data are added to the training set. The gray line shows the coefficient used to generate the map. Rather than monotonically converging to the true value, the estimated coefficient fluctuates in both magnitude and sign as data are added. This was common for all coefficients. We found that, on average, 65% of the pairwise coefficients flipped signs as data was added to our model.

These observations were robust to the choice of epistatic model and regression method. We used the Hadamard epistatic model with lasso regression for the results shown, but obtained identical results for all combinations of the Hadamard and biochemical epistatic models with ordinary, lasso, or ridge regression (see methods, Fig S2).

### Predictive epistatic models cannot be extracted from experimental genotype-phenotype maps

We next asked whether experimental genotype-phenotype maps exhibited similar bias in their regressed epistatic coefficients. We analyzed 12 experimentally characterized genotype-phenotype maps (Table 1). All maps contained all combinations of *L* mutations, ranging in size from 32 to 128 genotypes. The maps consisted of very different classes of genotypes: collections of point mutations within a single gene, scattered genomic point mutations, or alternate alleles of genes in a metabolic network. The measured phenotypes are also diverse: competitive fitness, binding affinity, and parameters like growth rate and sporulation efficiency.

**Table 1:**
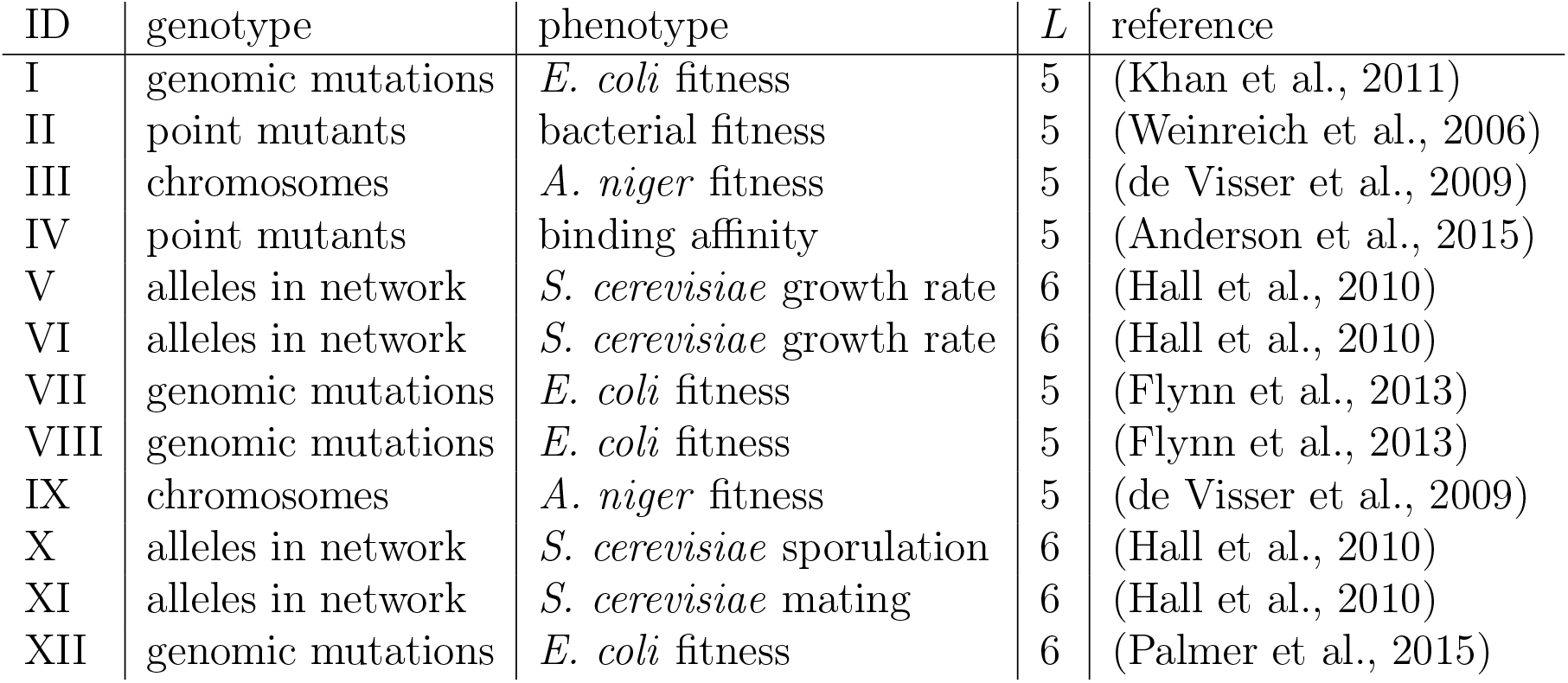
Published experimental genotype-phenotype maps.

We started by linearizing the experimental genotype-phenotype maps (see methods) (Sailer and Harms, 2017a; Otwinowski et al., 2018). We then dissected each map into linear epistatic coefficients. Because all genotype-phenotype pairs in these maps have been measured, we have the same number of observations as epistatic coefficients. We can therefore decompose epistasis into linear coefficients using a matrix transformation (Heckendorn and Whitley, 1999; Poelwijk et al., 2016), avoiding complications arising from regression. We found that all of these maps exhibited statistically significant pairwise and high-order epistatic interactions. Epistasis contributed from 6% to 79% of the variation in these maps (Fig 2A).

**Figure 2:**
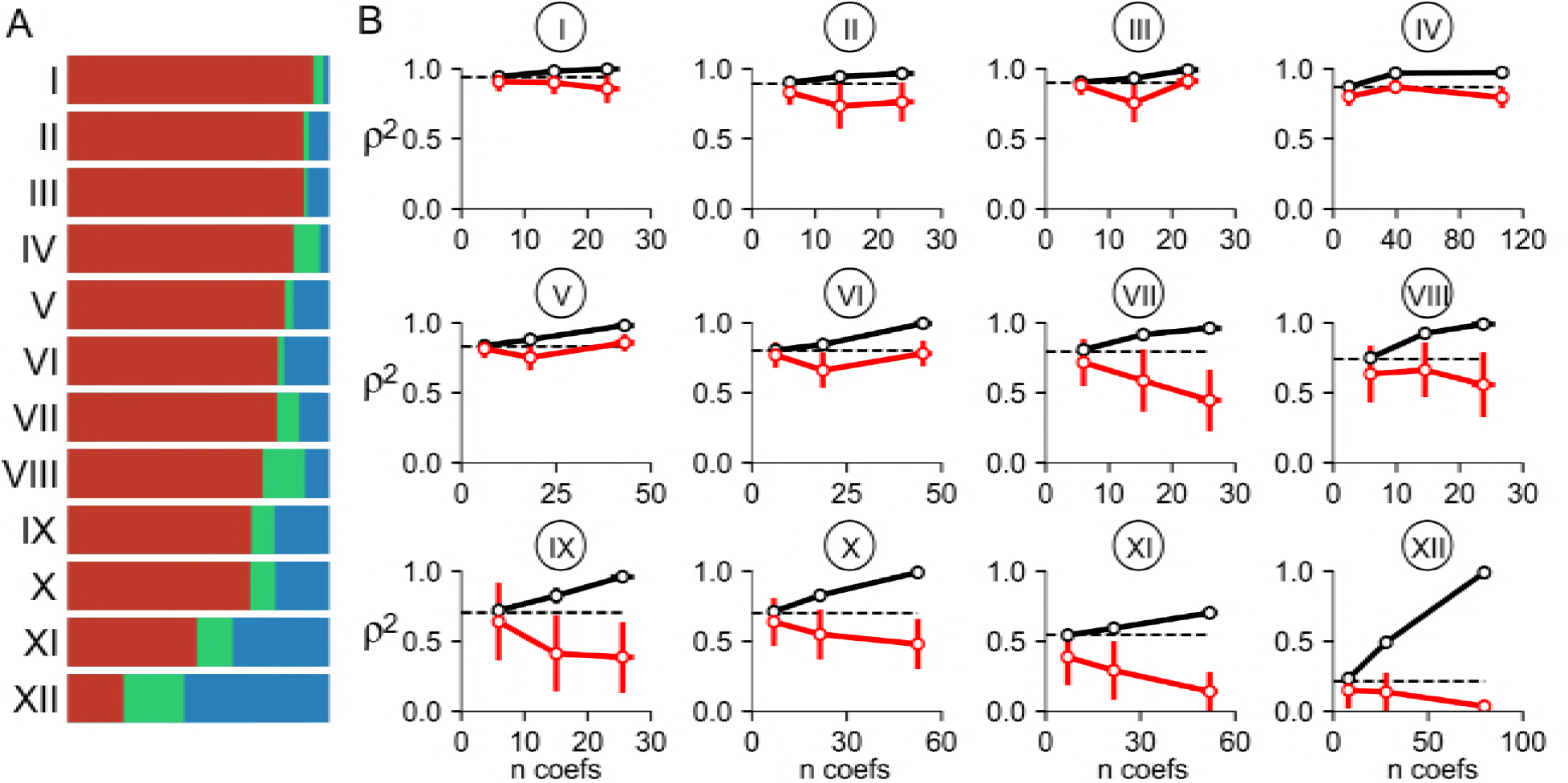
Predictive epistatic coefficients cannot be resolved from experimental genotype-phenotype maps. A) Bars show the fraction of variation in phenotype explained by additive effects (red), pairwise epistasis (green), or any order of high-order epistasis (blue). Each bar is for one of the twelve experimental genotype-phenotype maps. B) Each sub-panel shows 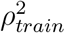 (black) and 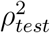 (red) for the map indicated above the graph as epistatic orders are added to the model. The x-axis is the number of parameters used in the fit. Points are, from left to right: additive, pairwise, and high-order epistasis. Points and lines indicate the mean of 1,000 pseudoreplicate samples. Error bars are standard deviation of pseudoreplicate results. The dashed lines indicate the fraction of the variation in the map explained by the additive model. These fits used the Hadamard model with lasso regression. See Fig S3 for other epistatic models and regression strategies.

We next probed our ability to extract predictive epistatic coefficients from the linearized maps. We created a training set consisting of 80% of the genotype-phenotype pairs in each map, regressed models against this set of observations, and then predicted the phenotypes of the remaining 20% of the genotypes. As above, we fit the additive, pairwise and high-order models. We then repeated this for 1,000 pseudo-replicate training and test sets on each map.

As with our simulations, we found we could not reliably extract predictive epistatic coefficients (Fig 2B). In 11 of 12 maps, the additive model performed better than any other model. In seven of the twelve maps (I, VII, VIII, IX, X, XI, and XII), 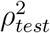 consistently decreased with each addition of epistatic coefficients. In four of the maps (II, III, V, and VI) the addition of pairwise epistasis led to a drop in 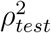 that was partially offset by the addition of high-order coefficients. Ultimately, however, the high-order model did no better than the additive model in these maps. Map IV was the the only map in which adding epistatic coefficients had any positive effect: the addition of pairwise epistasis led to a small increase in 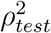 (from 0.80 to 0.87). This is achieved, however, by increasing the number of fit parameters from 10 to 40, implying that each epistatic coefficient contributed very little to the overall model. As with the simulated maps, these observations were robust to the choice of epistatic model and regression strategy (Fig S3).

### Experimental epistatic coefficients cannot be distinguished from a random model

These results indicate that predictive, linear epistatic coefficients cannot be estimated by regression in these genotype-phenotype maps. We must characterize essentially every phenotype in a genotype-phenotype map to resolve the epistatic coefficients that describe the map. But, if we have measured every phenotype, there are no more phenotypes to predict. One might conclude that understanding epistasis requires measuring *every* genotype-phenotype pair in a map.

Given the effort required to measure every phenotype, we posed another question: is it worth exhaustively characterizing a map just to extract epistatic coefficients? Or, put differently, are the epistatic coefficients one can decompose from a complete map informative? We approached this question by comparing the epistatic coefficients extracted from an experimental genotype-phenotype map to those extracted from a null model. Our null model was a random map: we generated phenotypes with an additive model and then perturbed each phenotype by a random value drawn from a normal distribution centered at zero. This is an appropriate null model because the generating model has no mechanistic interactions at all; any correlations between mutations arise from noise. Such a map consists entirely of “statistical” epistasis (Cordell, 2002).

We decomposed the epistasis in Map VIII using all 32 measured phenotypes and compared the resulting epistasis to our null model. Fig 3A-C shows the epistasis extracted from the experimental map. In this map 26% of the variation in phenotype is due to epistasis (Fig 3B). The residuals between the additive model and the observed phenotypes are normally distributed (Fig 3B). When we decompose the epistasis, we find that pairwise coefficients capture 16.2% and high-order coefficients capture 9.2% of the variation in phenotype.

**Figure 3:**
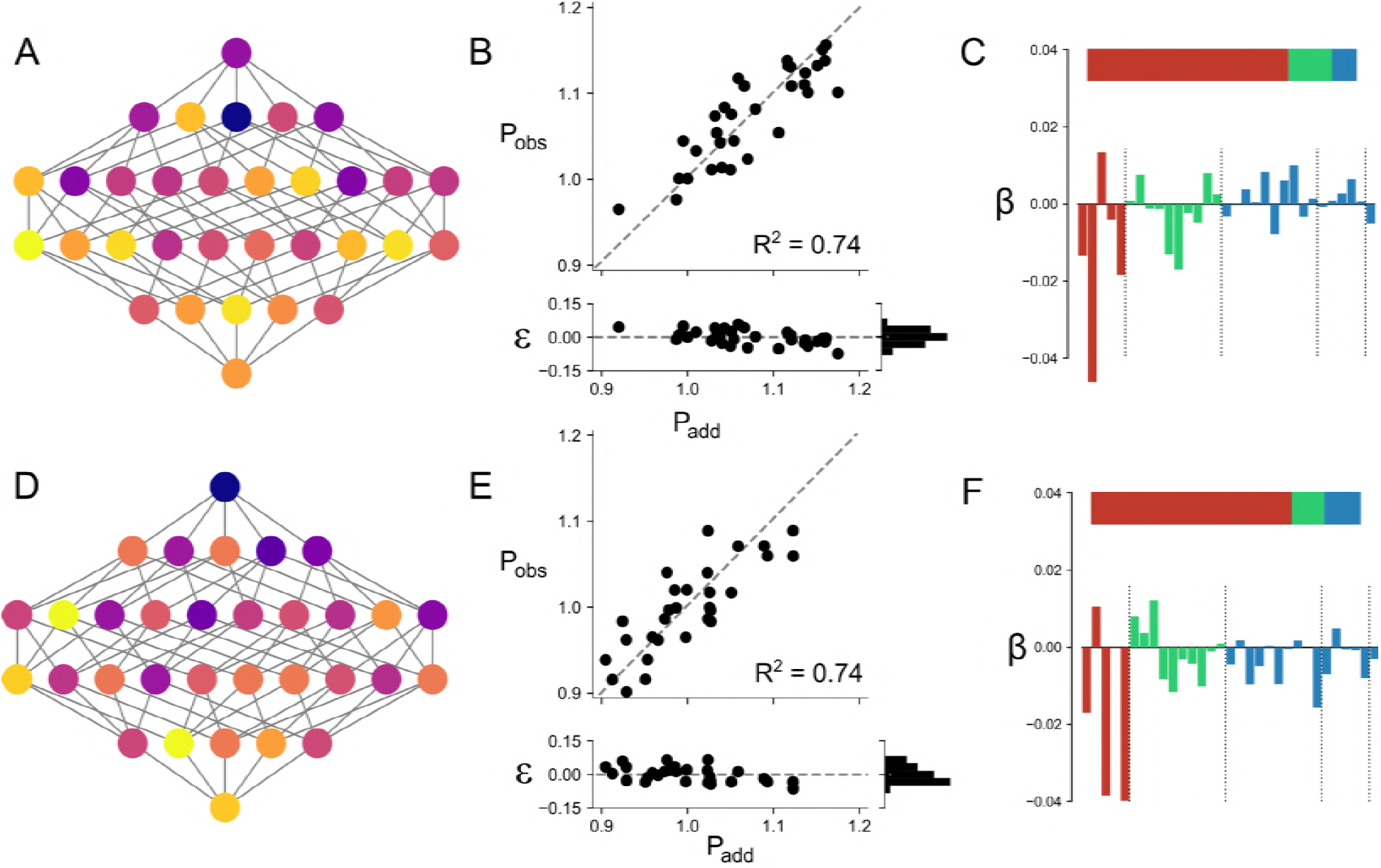
Experimental maps resemble random maps. Panels A and D show genotype-phenotype maps. Each node is a genotype; each edge is a single point mutation. Colors indicate quantitative phenotype. Panels B and E show the correlation between the observed phenotypes and the additive model, with fit residuals shown below the plot. Panels C and F indicate the magnitude of epistasis in each map as in Fig 2A (top subpanel) and the values of all model coefficients (bottom subpanel). Colors indicate additive components (red), pairwise components (green), and high-order components (blue). Each bar shows the value of a single model coefficient: the red bars correspond to the 5 additive coefficients, the green bars to the 10 pairwise coefficients, and the blue bars to the 17 high-order coefficients. Panels A-C are for experimental map VIII; panels D-F are for a simulated map with random epistasis.

We next constructed our null map. We generated a collection of random additive coefficients and calculated *P_add_* for each genotype. We then added random perturbations to each phenotype, drawn from a normal distribution with a mean of 0 and a standard deviation selected to yield a total magnitude of epistasis similar to the experimental map. This sampling procedure gave the *P_obs_* vs. *P_add_* curve shown in Fig 3E. As with the experimental map, epistasis accounted for 26% of the total variation in the map. We then decomposed this random epistasis with a high-order epistasis model (Fig 3F).

The overall structure of epistasis is indistinguishable between the experimental and a random map, even though the values of the specific epistatic coefficients are different (Fig 3C vs. F). If we generate many random maps—effectively, sampling over the possible configurations of epistatic coefficients that arise from a random variation in phenotype—we cannot distinguish the experimental map from among the decoys (Fig S4). This suggests that the linear epistatic coefficients extracted from this map should be viewed as decompositions of random noise, unless this can be shown otherwise.

### Using an additive model to treat epistasis

Our results speak against decomposing epistasis into collections of linear interaction terms. So how should we treat epistasis? We will touch on nonlinear treatments in the discussion, but before doing so, we will explore our top-performing epistasis model from above: the additive model.

The additive model treats epistasis as residual variation not explicitly accounted for by the model. If we measure the phenotypes of a set of combinatorial genotypes, we observe the effect of each mutation in a large number of genetic backgrounds (Fig 4A). We can describe the effect of mutation *i* with two numbers, its average effect 〈*β_i_*〉 and the variance of its effect 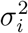. This same logic applies at the level of whole genotypes. If we have linearized the genotype-phenotype map (Szendro et al., 2013; Sailer and Harms, 2017a; Otwinowski et al., 2018), the residuals between 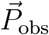 and 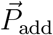 will be normally distributed (Fig 3B). As a result, the phenotype of a genotype *g* is given by:

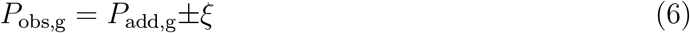

where ξ is the standard deviation of the residuals between 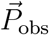 and 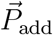. This is the basic definition of epistasis given by Fisher (Fisher, 1918), applied across the whole map.

**Figure 4:**
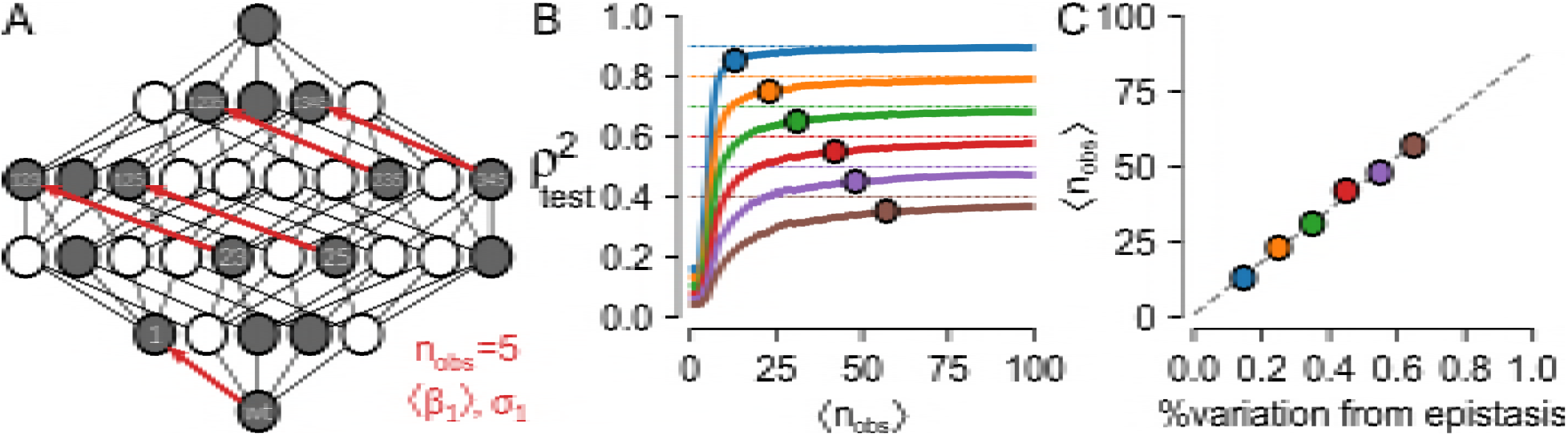
Epistasis as uncertainty. A) A partially characterized map. Circles represent genotypes, some of which have been measured (filled), some of which have not (unfilled). Lines represent single point mutations. Given these observations, we measure the effect of mutation 1 in five different backgrounds (red arrows) and can thus calculate the mean and variance in its effect across the map (〈*β*_1_〉 and *σ*_1_). B) 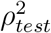 versus the average number of times each mutation is seen in randomly sampled genotype-phenotype maps with epistasis responsible for 10% (blue) to 60% (brown) of the variation in the maps. Points indicate where 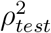 is within 5% of the maximum predictive power of the additive model. C) A calibration curve indicating how many times, on average, one must observe each mutation in map to resolve the additive coefficients in a map with different fractions of epistasis.

This view is particularly useful for predicting unmeasured phenotypes. First: it means each predicted phenotype has a known, normally distributed uncertainty. Even if a large amount of variation remains unexplained by the additive model, it is safely partitioned into a random normal distribution. Put another way, *ξ* acts as a prediction interval. Second: because the additive model has few terms, we can train it using a very small amount of data.

Following this line of reasoning, we asked how many phenotypes we would have to measure to construct a maximally predictive additive model. We constructed additive maps with different alphabet sizes (ranging from 2 to 5) and numbers of mutations (ranging from 6 to 8). We then injected random epistasis ranging in magnitude from 10% to 60% of the variation in the phenotype. We simulated experiments where we measured one random genotype at a time, added it to our observations, and predicted the phenotypes of the remaining genotypes. We then plotted 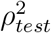 as a function of the average number of times we saw each individual mutation across all genetic backgrounds (〈*n_obs_*〉).

When plotted as a function of 〈*n_obs_*〉, 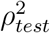 rapidly rises and then saturates at the magnitude of the epistasis in the map, independent of alphabet size and number of mutations (Fig 4B). We next asked, as a function of the magnitude of the epistasis in the map, when our predictions would be within 0.05 of the best achievable 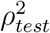. This is indicated by the points on Fig 4B. We plotted these values as a function of the magnitude of the epistasis in the map. This reveals a linear relationship between the average number of times we need to see each mutation and the total epistasis in the map (Fig 4C).

We set out to test this approach using a partially sampled, experimental genotype-phenotype map characterizing the binding specificity of dCas9 to 23-base-pair oligonucleotides (Fig 5A). The published experiment sampled 59, 394 of the 7 × 10^13^ (4^23^) possible oligonucleotides. Although all bases were sampled at all positions, there was significant bias towards a specific base at each position in the library (Fig 5A). The map exhibited a highly non-linear relationship between 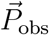 and 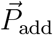 (Fig 5B), so we linearized the map with a 5th-order spline (Eq. 4), yielding normal residuals between 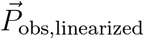 and 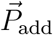 (Fig 5C). We then assessed the predictive power of the map: we added genotypes individually to a training set and evaluated our ability to predict the test set. We found that we were able to fit a model using ≈ 4, 000 genotypes to predict the remaining ≈ 55, 000 measurements. Because of the biased sampling of genotypes in the map, it took 4, 000 genotypes to observe each individual mutation a sufficient number of times to resolve the additive effects of all mutations (Fig 5D). Our prediction curve saturated after we had seen each mutation at least 39 times. This is in good agreement with our calibration curve on simulated data, which indicated we would need to observe each mutation an average of 40 times (with random sampling) to saturate an additive model in which epistasis was responsible for 38% of the variation in the map.

**Figure 5:**
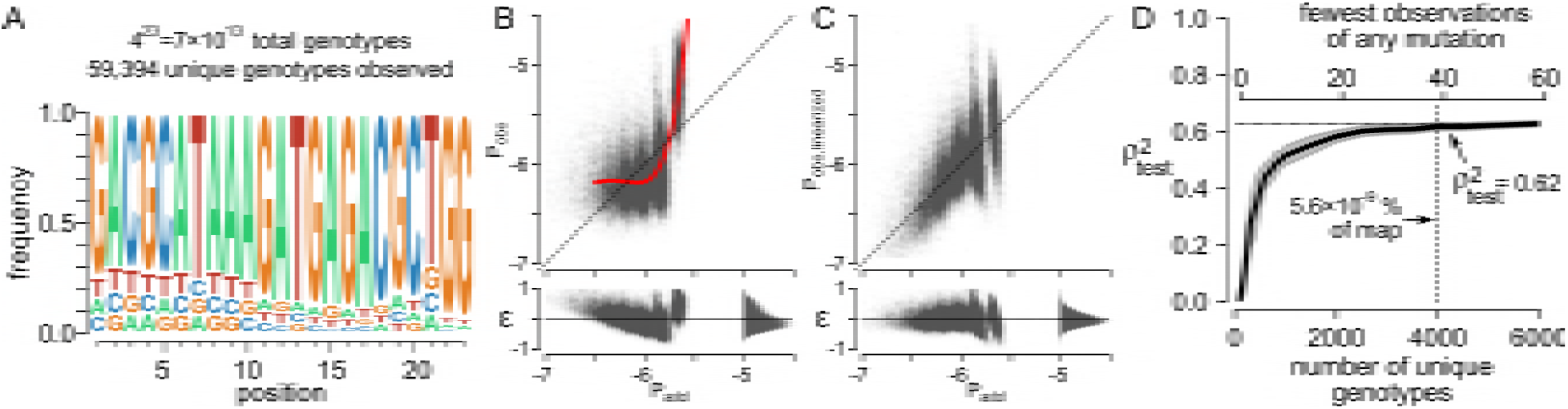
A predictive, additive model can be trained on a large genotype-phenotype map. A) Summary of the genotype-phenotype map reported in (Boyle et al., 2017). Map consists of 23 sites, each with four bases with the frequency at each site shown in the sequence logo. The total map has 7 × 10^13^ genotypes; the publication reports measured phenotypes for 59, 394 genotypes. B) Raw *P_obs_* vs. *P_add_* plot for the map. Each point is a genotype. The fit residuals are shown below the main plot. We fit an 5th-order spline to linearize the map (red curve). C) The linearized form of the map, with epistasis removed using the spline shown in panel B. D) A predictive model can be trained using ≈ 4, 000 genotypes. The bottom x-axis shows the number of unique genotypes used to train the model (sampled randomly); the top x-axis shows the fewest number of times any mutation was seen in that sample given the bias in the frequencies of the input mutations. 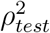 was measured against the remaining 50, 000+ genotypes not used to train the model.

The predictive power of this model is quite good considering its simplicity: we are able to predict any phenotype to ±38% given we only sampled one, one-billionth of a percent of the map. Extensive epistasis remains, but it follows a normal distribution with a known standard deviation. While there are certainly more sophisticated models, an additive model provides significant predictive power for this map.

## Discussion

Our results suggest that a linear model should not be used to extract pairwise and multi-way interactions between mutations in a genotype-phenotype map. Regressed epistatic coefficients are biased (Fig 1B), unstable to the addition of new data (Fig 1C), and not useful for predicting unmeasured phenotypes (Fig 2A). Far from being an anomaly, this appears to be a shared feature of a collection of a dozen high-precision, combinatorially-complete genotype-phenotype maps (Fig 2B). Further, we can generate epistatic coefficients very similar to those observed in these maps using a simulated map in which we added random, normally distributed noise to each phenotype (Fig 3). This argues for viewing epistatic coefficients as uninterpretable decompositions of random variation, unless shown otherwise.

Viewed mechanistically, this is unsurprising. The epistatic models under investigation assume linearity, but biology is nonlinear (Bershtein et al., 2006; Lehner, 2011; Chou et al., 2011; Tokuriki et al., 2012; Schenk et al., 2013; Sailer and Harms, 2017a; Otwinowski et al., 2018). There is no reason to believe that a linear model will capture a complicated non-linear system in a predictive and interpretable way. For example, we showed recently that we could generate high-order epistasis using a toy thermodynamic model of proteins with only explicit pairwise interactions (Sailer and Harms, 2017b). The epistasis arise because mutations have a nonlinear effect on the relative populations of individual protein conformations. As a result, epistatic coefficients cannot be interpreted mechanistically—they are purely “statistical” (Cordell, 2002).

Further, our results indicate that the signs and magnitudes of specific epistatic interactions extracted from genotype-phenotype maps have no universal meaning. For example, in Fig 1C, the selected pairwise coefficient flips between positive, zero, and negative. If different genotypes of the map are characterized, we obtain different values for the pairwise coefficient, and thus a different interpretation for the effect of epistasis on the phenotype.

### Treating epistasis with an additive model

A simple way to treat epistasis is as the residual variation after fitting an additive model (Eq. 6). Despite its simplicity, this is a useful perspective. It can be used to predict unmeasured phenotypes in a genotype-phenotype map with known uncertainty. This is because deviation from the additive model is determined by the magnitude of epistasis in the map (Fig 4A). Further, the simplicity of the model means we can characterize an extremely sparse sample of combinations of mutations across a genotype-phenotype map and still predict missing phenotypes (Fig 4C).

This suggests that sparsely sampling combinatorial genotypes, rather than aiming to exhaustively characterize point mutants, may be a powerful way to understand and predict genotype-phenotype maps. As long as each mutation is seen across a sufficiently large number of genetic backgrounds, we can resolve its average effect across a volume of the genotype-phenotype map. In contrast, exhaustively sampling point mutations in a single background— such as a deep mutational scan—will yield mutational effects specific to whatever genetic background is used. Epistasis is not averaged out, meaning such coefficients should not provide high predictive power when mutations are combined.

Interpretation of epistasis as a prediction interval only holds when the fit residuals are normally distributed about zero. Curvature between 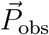 and 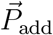 will lead to non-normal residuals and, thus, a distorted picture of the uncertainty (Sailer and Harms, 2017a; Otwinowski et al., 2018). Multiple methods exist for linearizing genotype-phenotype maps, including taking the log of phenotypes (Cordell, 2002), power transforms (Sailer and Harms, 2017a), splines (Otwinowski et al., 2018), and even mechanistic models (Schenk et al., 2013; Otwinowski, 2018). This is one area for improvement, as better global models will decrease the amount of variation that must be explained by the additive model. For example, in our analysis of the dCas9 binding specificity, there is still structure in the residuals, with some clustering along *P_add_* despite linearization using an 5th-order spline (Fig 5C). A model that captures such variation could improve predictive power. Further improvement of global models will thus be an important area of investigation.

### Moving away from linear models

We see three promising ways forward. The first is to view epistasis in terms of its consequences for evolutionary trajectories. This includes metrics like the number of accessible trajectories, number of fitness peaks, and summary statistics such as the roughness to slope ratio (Szendro et al., 2013; Ferretti et al., 2018; Crona et al.). These metrics generally do not allow prediction of unmeasured phenotypes nor mechanistic understanding of the map, but can provide useful insights into evolutionary trajectories and outcomes without the poor behavior observed in linear epistasis models.

The second is to use non-biological, nonlinear models to extract information from each map. These include tools such as Potts models (Figliuzzi et al., 2016; Hopf et al., 2017), variational auto encoders (Riesselman et al., 2017; Sinai et al., 2017), and neural networks (Wang et al., 2017; Ma et al., 2018). Such approaches can yield predictive models of genotype-phenotype maps, and will no doubt continue to grow in popularity and sophistication. One downside to these models is a requirement for massive amounts of training data—which may not always be feasible, even in the modern high-throughput era. Further, it may be difficult to link such models to an underlying biological mechanism.

The third is to attempt to model the underlying mechanistic process that leads to the map (Tokuriki et al., 2012; Schenk et al., 2013; Otwinowski, 2018; Dutta et al., 2018). Rather than taking a “top-down” approach, in which one dissects epistasis into statistical interactions that are hopefully meaningful, one can instead take a “bottom-up” approach, in which one calculates phenotypes from genotypes using a mechanistic biological model. This model can then be trained against measured phenotypes. This provides a predictive model for unmeasured phenotypes, as well as providing mechanistic insight into map between genotype and phenotype. A good example is that of Schenk et al, who dissected a genotype-phenotype map by explicitly modeling the effect of each mutation on protein stability and enzymatic activity (Schenk et al., 2013). This model captured extensive variation in the map that could not be described with a linear model, while also providing mechanistic insight into the protein under investigation.

### Conclusion

Epistasis was described by Fisher as residual variation left over after fitting an additive model (Fisher, 1918). While it may sometimes be productive to separate these residuals into specific statistical coefficients, a better approach is to build better model. In our view, the long-term goal should not be interpreting epistatic interactions between mutations; rather, the long-term goal should be building mechanistic models that fit experimental observations and, ultimately, make epistasis disappear.

## Acknowledgements

We would like to thank members of the Harms lab for helpful discussions and comments. Work was supported by start up funds from the University of Oregon (ZRS). MJH is a Pew Scholar in the Biomedical Sciences, supported by The Pew Charitable Trusts.

